# How much pollen do solitary bee larvae consume? : Or establishing realistic exposure estimates of solitary bee larvae via pollen for the use in risk assessment

**DOI:** 10.1101/2021.05.10.443351

**Authors:** Tobias Pamminger, Christof Schneider, Raffael Maas, Matthias Bergtold

**Affiliations:** BASF SE, Speyererstraße 2, 67117 Limburgerhof, Germany; Bayer AG, CropScience division, Alfred-Nobel-Straße 50, 40789 Monheim

## Abstract

Bees foraging in agricultural habitats can be exposed to plant protection products. In order to limit the risk of adverse events to occur a robust risk assessment is needed, which requires reliable estimates for the expected exposure. Especially the exposure pathways to developing solitary bees are not well described and in the currently proposed form rely on limited information. To address this topic, we used a published data set on the volume of pollen solitary bees provide for their larvae to build two scaling models predicting the amount of protein and pollen developing solitary bees need based on adult body weight. We test our models using both literature and experimental data, which both support the validity of the presented models. Using scaling models in the bee risk assessment could complement existing risk assessment approaches, facilitate the further development of accurate risk characterization for solitary bees and ultimately will help to protect them during their foraging activity in agricultural settings.

## Introduction

In recent years there is growing concern that non-managed bee populations are in decline, potentially compromising pollination security in agricultural and non-agricultural landscapes (Gallai, Salles et al. 2009, Grab, Branstetter et al. 2019). While numerous drivers are likely associated with this trend, the exposure of bees to plant protection products (PPP) could be one of them (Goulson, Nicholls et al. 2015). Consequently, bees need to be protected from potentially adverse events and risk assessment (RA) schemes are in place for the registration of PPPs prior to their placement on the market (EFSA 2013, USEPA 2014, AAVM 2015).

Historically, due to the technical and logistic difficulties of testing non-commercially raised species, the established pollinator RA procedures (e g. EFSA and EPA) have relied on the honeybee (*Apis mellifera*) as surrogate species. In contrast to the majority of bees *A. mellifera* is highly social (Danforth, Minckley et al. 2019), which can have important consequences for the RA in particular for the expected exposure of different developmental stages and scenarios as they do not directly provide pollen to their offspring (Boyle, Pitts-Singer et al. 2019). While the currently implemented *A. mellifera* centered pollinator RA schemes are likely protective for solitary bees (SB) as well (Boyle, Pitts-Singer et al. 2019, Thompson and Pamminger 2019) it is unclear if this also extends to exposure routs not directly addressed in current pollinator risk assessments (Boyle, Pitts-Singer et al. 2019). One of these alternative exposure routes is related to developing SB, which in contrast to honeybees often feed on a single provision of unprocessed, and potentially PPP contaminated pollen mixed with varying degrees of nectar (Boyle, Pitts-Singer et al. 2019). While nectar can be contaminated with PPP residues as well recent evidence suggest that the main driver of PPP residue in SB larvae provisions is likely pollen (Kyriakopoulou, Kandris et al. 2017). Consequently, an accurate estimate of SB larvae pollen provisions is critical to evaluate the potential risk SB larvae face from PPP residues during this time. However, the currently proposed estimates for pollen consumption of SB larvae relies mainly on limited information from a restricted number of species making their accuracy, robustness and generalizability uncertain (Ladurner, Maccagnani et al. 1999, Bosch and Vicens 2002, EFSA 2012).

In this study, based on a published data set on the pollen volume provided to SB larvae (Müller, Diener et al. 2006), we develop generalized and RA compatible scaling model to directly predict the pollen provisions [mg] of SB based on adult bee body weight and test its predictions using both published and experimentally generated data.

## Material, Methods & Results

All statistics and visualizations were conducted in the R statistical environment (R 2013) v 4.0.3 using Rstudio Version 1.4.1103 (RStudio 2020). We used the following packages: ggplot2 (Hadley 2016), ggpubr (Kassambara 2020), dplyr (Hadley, Romain et al. 2021), ggrepel (Slowikowski, Schep et al. 2016) and tidyverse (Wickham, Averick et al. 2019).

### DATA

We use a published data set (Müller, Diener et al. 2006), describing the association between adult SB species’ dry weight (mean dw [mg]) and the volume [mm^3^] of pollen provisioned. The dataset includes information for female specimens of 14 SB species from Europe and in four cases measurements for male as well (total N = 18). We replicate the data by fitting a liner model (LM) with a random intercept to the log_10_ transformed bee weight (predictor variable) and pollen provision volume (response variable) following the authors initial analysis (Müller, Diener et al. 2006). We confirm their findings demonstrating clear linear association between the two log_10_ transformed variables (LM: F = 46.41; df = 16; R^2^ = 0.74, P < 0.001; Equation 1; Fig. 1).

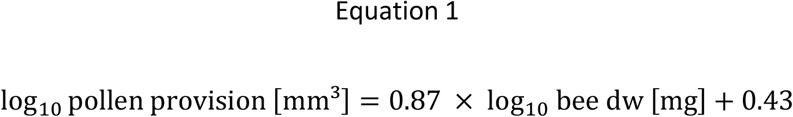

**Figure 1:**
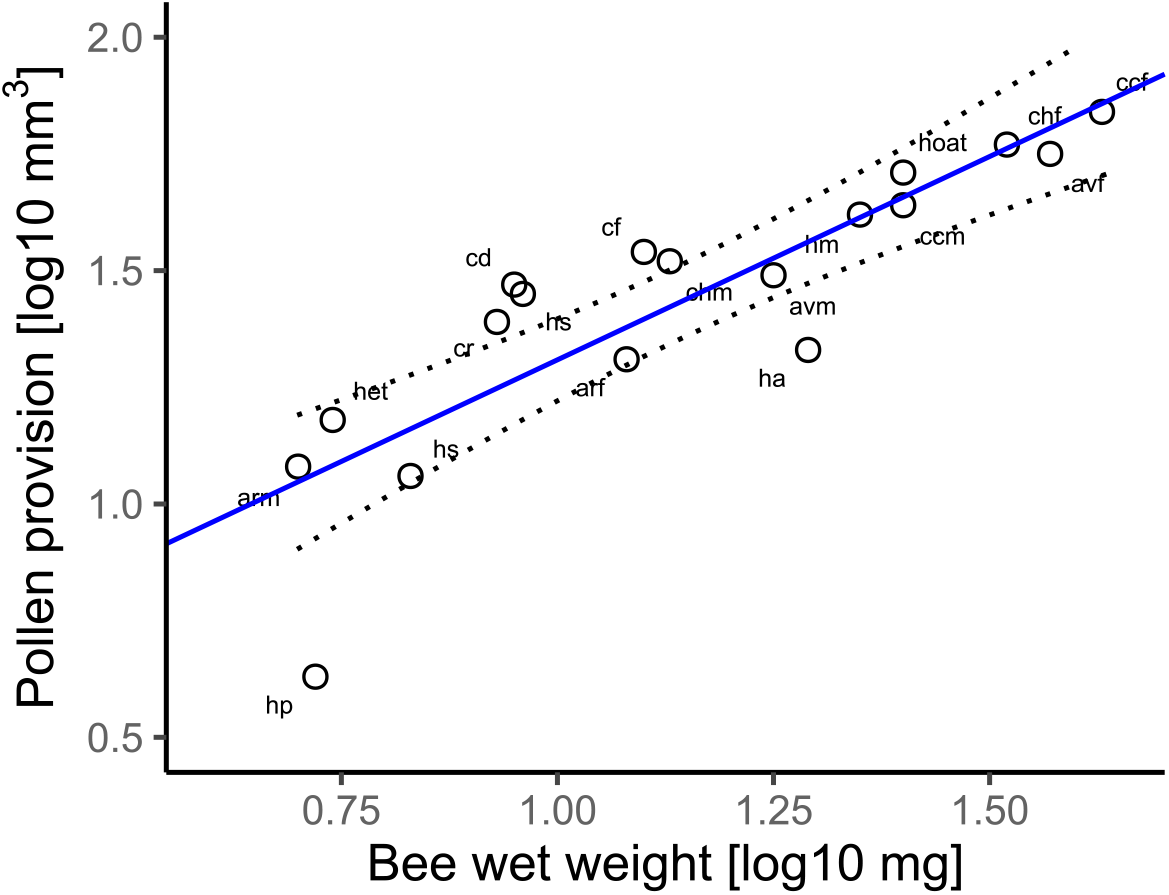
showing the relationship between bee dry weight in mg and the volume of pollen in mm^3^ provisioned for the developing larvae (LM: F = 46.41; df = 16; R^2^ = 0.74, P < 0.001; See Müller et. al 2006) and the associated 95%CI (dotted line). Arf = *Andrena ruficrus* (female); arm = *Andrena ruficrus* (male); avf = *Andrena vaga* (female); avm = *Andrena vaga* (male); ccf = *Colletes cunicularius* (female); ccm = *Colletes cunicularius* (male); cd = *Colletes daviesanus*; chf = *Colletes hederae* (female); chm = *Colletes hederae* (male); cf = *Chelostoma florisomne*; cr = *Chelostoma rapunculi*; ha = *Hoplitis adunca*; hm = *Hoplitis mocsaryi*; hoat = *Hoplitis tridentata*; het = *Heriades truncorum*; hos = *Hoplosmia spinulosa*; hp = *Hylaeus punctulatissimus*; Hs = Hylaeus signatus.

### PREDICTING THE CORRESPONDING PROTEIN PROVISION

Assumption that larvae protein requirements are an important driver determining the pollen provisioned volume we expected that accounting for the variation of protein content in host plant pollen would improve the models fit. We used host plant preferences of the SB species present in the data set (Müller, Diener et al. 2006, Westrich 2018) to determine the likely protein concentration [%] in the collected pollen provisions (Pamminger, Becker et al. 2019). We used the median plant genus estimates or family level information if genus level information was not available (see data set). In case of pollen generalist bee species (i e. bees collecting pollen from multiple plants genera) we used the median protein concentration of their reported host plant genera. Using the information, we calculate the expected volume of protein provided to SB larvae [mm^3^] and estimated the corresponding amount of protein [mg] using the reported mean protein density estimate of 1.37 [mg/mm^3^] (Erickson 2009). In order to see if this correction improved the scaling association, we fit a LM to the log_10_ protein provision [mg] and bee adult dry weight [mg]. The new model fit the data better indicated by the improved R^2^ values compared to the original model (LM: F = 166.5; df = 16; R^2^ = 0.91; P < 0.001; Figure 2).

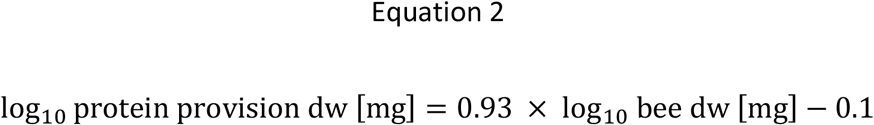

**Figure 2:**
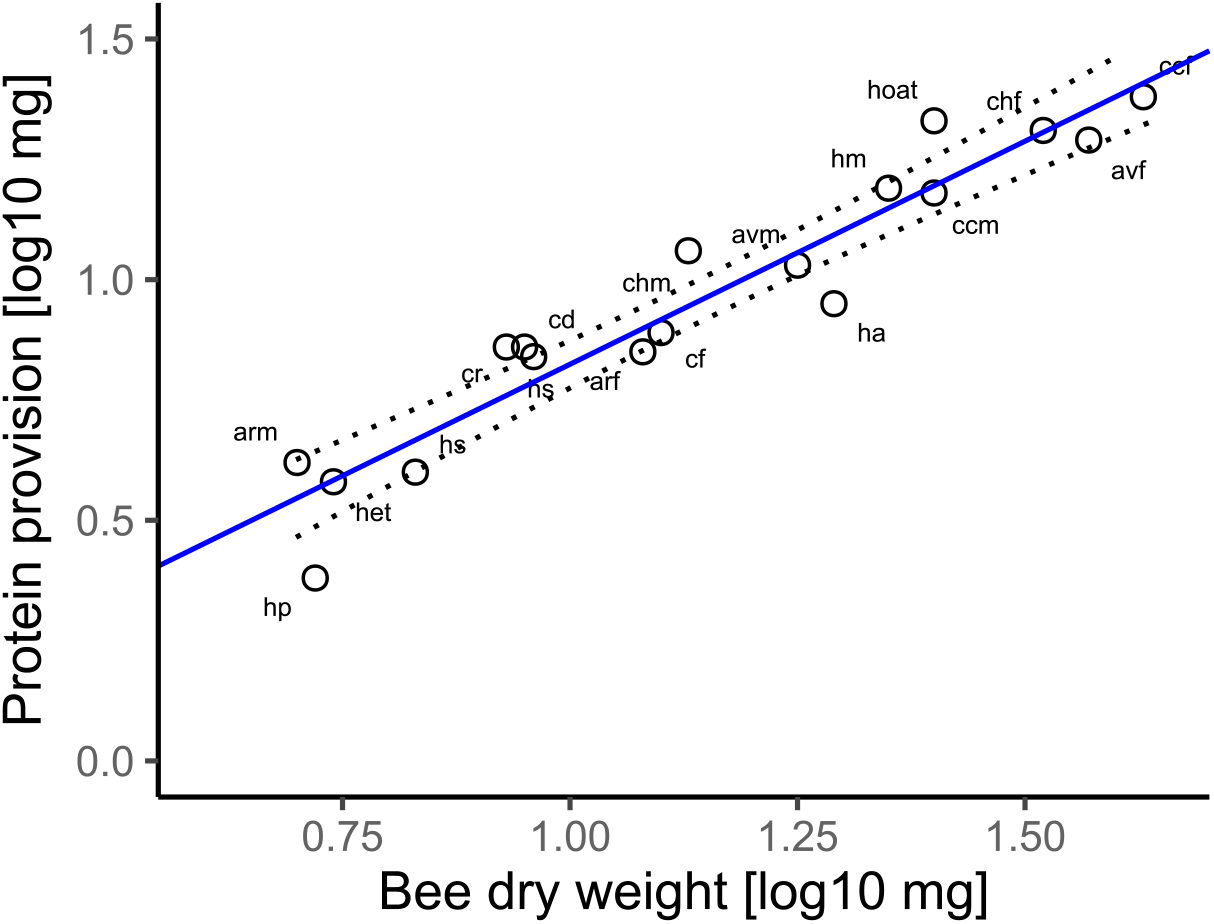
showing the relationship between bee dry weight in mg and expected protein provisioning in mg for the developing larvae (LM: F = 166.5; df = 16; R^2^ = 0.91; P < 0.001) and the associated 95%CI (dotted line). Arf = *Andrena ruficrus* (female); arm = *Andrena ruficrus* (male); avf = *Andrena vaga* (female); avm = *Andrena vaga* (male); ccf = *Colletes cunicularius* (female); ccm = *Colletes cunicularius* (male); cd = *Colletes daviesanus*; chf = *Colletes hederae* (female); chm = *Colletes hederae* (male); cf = *Chelostoma florisomne*; cr = *Chelostoma rapunculi*; ha = *Hoplitis adunca*; hm = *Hoplitis mocsaryi*; hoat = *Hoplitis tridentata*; het = *Heriades truncorum*; hos = *Hoplosmia spinulosa*; hp = *Hylaeus punctulatissimus*; Hs = *Hylaeus signatus*.

### PREDICTING POLLEN PROVISIONS

Because pollinator exposure assessment is based on the provisioned pollen and the expected or measured PPP residues within it we rescaled the protein prediction model (Equation 2), assuming a median pollen protein concentration of 29.1% (Pamminger, Becker et al. 2019), to directly predict the corresponding amount of pollen [mg] likely provisioned by a generalist SB based on their body weight.

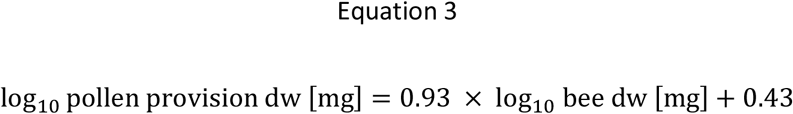

### TESTING THE MODEL PREDICTION

In a first step we tested the protein models’ predictions (equation 2) using published data on the weight for adult and protein needs for developing honey bee *Apis mellifera*. Based on the reported dry weight of an *A. mellifera* worker 25.52 mg (Kendall, Rader et al. 2019) the model predicts 15.89 (95% prediction confidence interval: 13.83-18.24)) mg dw protein are needed per bee. The amount of dw protein contained in an newly emerged adult worker (without intestinal tract) is about 16 mg (Hrassnigg and Crailsheim 2005), which is within the predicted 95%CI of the model supporting its validity (Figure S1).

In a second step we tested the validity of model 3 and predicted the expected pollen provision of a commercially available, regulatory relevant and pollen generalist (Westrich 2018) SB species (*Osmia bicornis*). Based on the reported mean body weight of male and females of 21.72 [mg dw] (Kendall, Rader et al. 2019) the model predicts an pollen provision of 46.98 (95% prediction confidence interval: 41.4 -53.21) mg of dw pollen. To test this prediction we set up a field experiment in June 2020 and released *O. bicornis* (total N = 430 males and 215 females supplier = WAB Mauerbienenzucht https://www.mauerbienen-shop.com/) in approximately equal proportions at three open agricultural location (L1-L3) and in one flight tunnel (L4; host plant *Phacelia spp*.) in south-west Germany (Limburgerhof). All locations were provided with artificial nest sites and the similar to the methods reported for Osmia field testing (Franke, Elston et al. 2021). After one initial week of acclimatization we checked the nest sites twice a week for newly built cells and removed the pollen provisions before the larvae had hatched and stored them in individual Eppendorf® tubes at -20°C till analysis. Over the duration of three weeks we collected a total of 161 pollen provisions and measured their wet- and dry weight as well as the corresponding amount of water-, sugar and pollen in mg following established methodology (Kapheim, Bernal et al. 2011). The pollen provisions were weighted (wet weight) to the closest 0.1 mg using a Mettler Toledo Laboratory Balance (XPR205DR) scale. In a next step the provisions were dried at 60°C over night and reweighted (dry weight). The water content was calculated as the difference between the two measurements. The dried pollen was suspended in 0.5-0.75 ml deionized water (depending on the provision size) and vortexed for 2 minutes to ensure the transition of all soluble sugars into solution. After centrifugation to a loose pellet at 6000 rpm for 1 min using a VWR Galaxy Mini Star centrifuge the dissolved sugars in the supernate were quantified using a hand-held refractometer (Bieno ®Vinum Refraktometer für Winzer und Mostereien) following the manuals instruction. The amount of dw pollen was calculated by subtracting the measured amount of sugar in mg from the pollen dry weight. The summary statistics for all locations are presented in Table 1. We find that *O. bicornis* provided 64.23 (95% prediction confidence interval: 47.86 -85.11) median mg dw pollen in agreement with the model predictions (overlapping 95%CI, Figure 3).

**Table 1:** Summarizing the results of the *O. bicornis* provision composition at the four sampled locations. For all locations we show wet and dry weight of the provisions as well as their water, sugar and pollen content. We present median and associated standard deviations.

| Location | N | wet weight [mg] |  | dry weight [mg] |  | water [ml] |  | sugar [mg] |  | pollen [mg] |  |
| --- | --- | --- | --- | --- | --- | --- | --- | --- | --- | --- | --- |
|  |  | median | SD | median | SD | median | SD | median | SD | median | SD |
| L1 | 51 | 168.3 | 76.5 | 132.3 | 60.11 | 37.2 | 18.39 | 74.2 | 35.46 | 57.9 | 27.3 |
| L2 | 27 | 201.4 | 73.6 | 171.4 | 60.17 | 30.0 | 14.51 | 78.0 | 28.43 | 92.1 | 37.4 |
| L3 | 32 | 161.4 | 94.3 | 132.1 | 76.57 | 28.9 | 18.78 | 58.7 | 42.91 | 55.5 | 40.4 |
| L4 (Tunnel) | 51 | 257.1 | 107.2 | 190.9 | 81.91 | 54.1 | 28.44 | 121.1 | 49.44 | 70.6 | 39.7 |
| Overall median | 161 | 184.9 |  | 151.8 |  | 33.6 |  | 76.1 |  | 64.2 |  |

**Figure 3:**
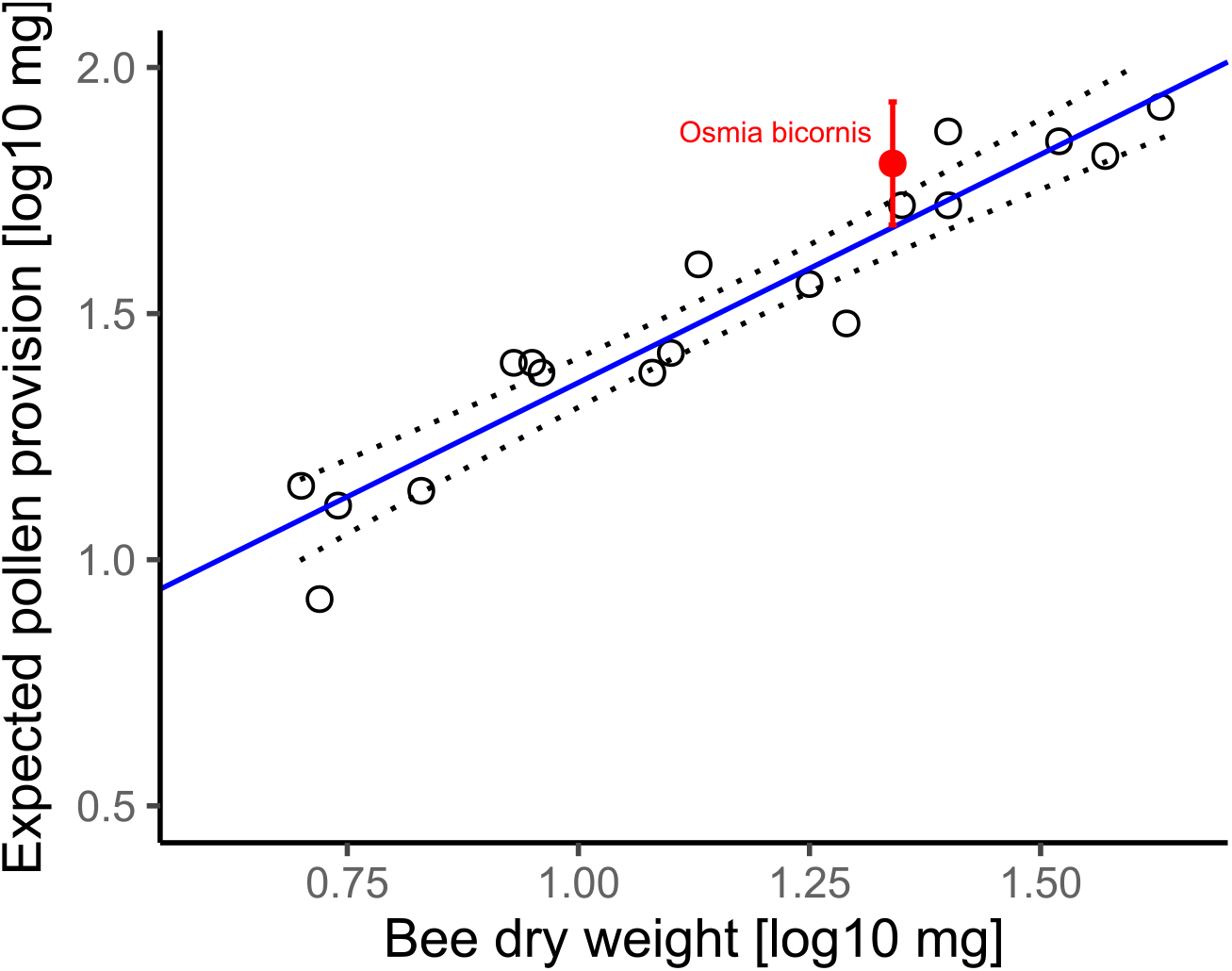
depicts the relationship between bee dry weight and the expected pollen provisioning for the developing larvae in mg (LM: F = 166.5; df = 16; R^2^ = 0.91, P < 0.001) and the associated 95%CI (dotted line). We compare these expectations to the measured pollen provisions of *O. bicornis in 2020* (in red; median_observed_ = 64.23 mg and associated 95% CI).

### FINAL MODEL

In a last step we updated the pollen provision prediction model by included the *O. bicornis* measurements (LM F = 161.9; df = 17; R^2^ = 0.91; P < 0.001).

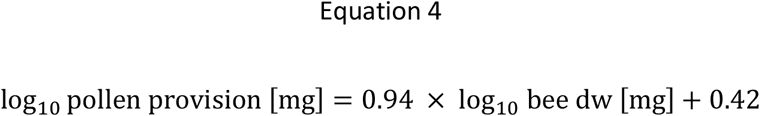

## Discussion

In this study we developed and tested two scaling models to predict the protein (equation 2) and pollen provisions (equation 3) of developing SB based on the corresponding adult dry weight for the use in pollinator risk assessment.

When looking at the protein prediction model (equation 2) we find that the linear model fits the log_10_ transformed data well (R^2^ = 0.91) across a wide weight range (4.6 – 43.1 mg dw) and can predict the approximate protein needs of developing *A. mellifera* reported in the literature (see results). This suggests that it might be possible that this equation applies to all bees independent of their social organization, because the observed relationship is likely driven by conserved protein needs of developing bees which in turn mainly depend on their size (larger bees need more proteins) and not their social organization. However, it is unlikely that this holds true for the presented pollen prediction model. In contrast to SB species, social bees pre-process pollen to varying degrees and often continuously provide it to their offspring during their larval stage (Gould and Gould 1988, Westrich 2018, Danforth, Minckley et al. 2019). Since it is in many cases unknown to which degree these alternate feeding patterns change the direct pollen exposure of developing social bees it is unlikely that the observed relation between SB bee size and larvae pollen needs can be directly extended to social bees without accounting for species or group specific differences.

SBs often provide their offspring with a single provision of unprocessed pollen of known host plant origin (Westrich 2018, Danforth, Minckley et al. 2019), which makes it possible to extrapolate their pollen needs directly from their protein requirements whenever the pollen protein concentration of the host plant(s) is known (Pamminger, Becker et al. 2019). In this case our pollen provision model is based on the median protein concentration found in the pollen of bee pollinated flowers which is likely a good approximation for pollen generalist bees. Using this model we were able to predict the pollen needs of the known pollen generalist *O. bicornis* (Westrich 2018) with good accuracy supporting the validity of the model (see figure 3). While the presented model seems well suited to predict the needs of pollen generalists it might be less accurate to predict the needs of pollen specialist bee species in particular ones preferring pollen with extremely high or low protein concentrations (Westrich 2018). In such cases it might be beneficial, similar to the procedure outline in this paper, to rescale the protein model (equation 2) considering the host plant preferences and the associated pollen protein concentrations.

When looking in more detail at the *O. bicornis* provisions sampled at the four locations, we find some variation in provision size and relative composition (Tab.1). Considering the low number of locations (tunnel N =1, field N = 3) we did not conduct a formal statistical, but rather describe the observed patterns qualitatively. Overall, the tunnel location (*Phacelia spp*. only) provided the largest provisions (median wet and dry weight) and contained the most sugar and water (nectar). In contrast the observed pollen provisions observed in the tunnel setting are within the observed values of the free flying locations. This could indicate that pollen provision might not be as dependent on host plant type and abundance compared to nectar (in both cases their supply can be expected not to be limited in the tunnel), but more independent observation is needed to draw definitive conclusions. When looking at the nectar content we see that in the tunnel the provisions are larger mostly due to their sugar and water (nectar) content. This is in line with recent findings showing that if given the choice *O. bicornis* bees will favor carbohydrates over pollen if given the choice (Austin and Gilbert 2021). One clear limitation of our sampling procedure is that we do not know the sex of the developing larvae. Since it is known that the size of the provisions can be sex dependent in the *Osmia* genus (Bosch and Vicens 2002) and that *O. bicornis* sexes vary in size (Kendall, Rader et al. 2019) site dependent variation in sex ratios might account for some of the location specific variation in provision size. However, it is likely that the presented model can be applied to both sexes as the original data set encompassed both male and female measurements for four species and no obvious sex specific deviations were observed in protein or pollen need (see Figure 1 and 2).

Scaling approaches based on body weight can be used to estimate both hazard and exposure parameters in a range of organisms and some are currently utilized in ecological risk assessment schemes (Davidson, Parker et al. 1986, Urban and Urban 1986, Mineau, Collins et al. 1996, Mineau, Baril et al. 2001, EFSA 2009, Pamminger 2021). Such methods offer a clear alternative to experimental investigations in cases where such approaches are not feasible (e g. number of species) or desirable (e.g vertebrate testing). Similarly our models could be directly used to extend the currently implemented pollinator RA schemes to better cover SB specific exposure scenarios, which in turn would allow a more accurate risk evaluation for SB foraging in agricultural habitats and ultimately help to better protect them.

## Acknowledgment

We would like to thank Dr. Magdalena Mair for carefully reading of an earlier version of the manuscript and for providing insightful comments and a non-regulatory perspective on the topic.

---

**Figure S1:**
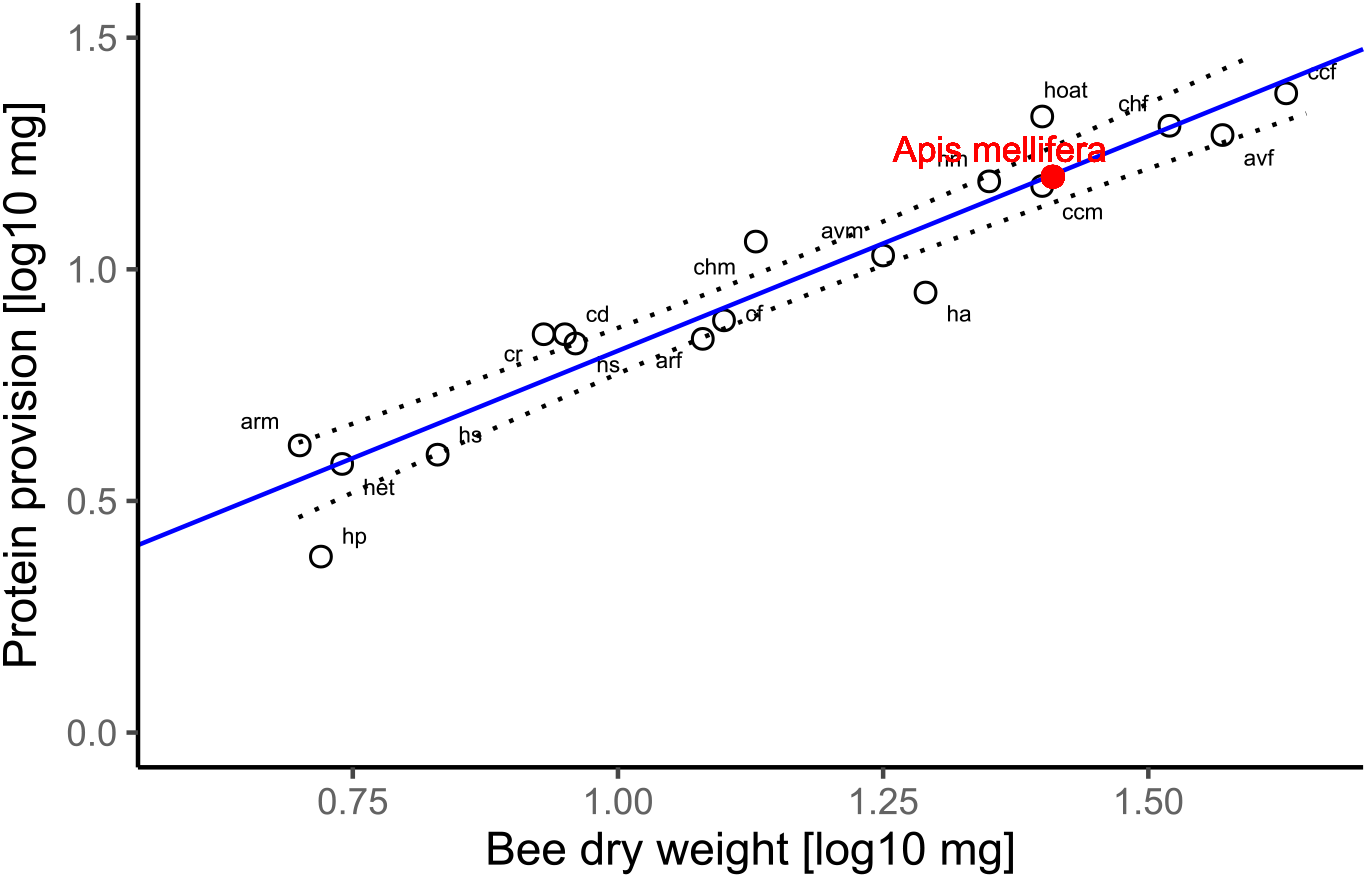
depicts the relationship between bee dry weight and the expected protein provisioning in mg for the developing larvae (LM: F = 166.5; df = 16; R^2^ = 0.91; P < 0.001) and the associated 95%CI (dotted line). In red we have plotted the reported protein provision for *A. mellifera*, which was not part of the SB data set published by Müller et al. 2006. Arf = *Andrena ruficrus* (female); arm = *Andrena ruficrus* (male); avf = *Andrena vaga* (female); avm = *Andrena vaga* (male); ccf = *Colletes cunicularius* (female); ccm = *Colletes cunicularius* (male); cd = *Colletes daviesanus*; chf = *Colletes hederae* (female); chm = *Colletes hederae* (male); cf = *Chelostoma florisomne*; cr = *Chelostoma rapunculi*; ha = *Hoplitis adunca*; hm = *Hoplitis mocsaryi*; hoat = *Hoplitis tridentata*; het = *Heriades truncorum*; hos = *Hoplosmia spinulosa*; hp = *Hylaeus punctulatissimus*; Hs = *Hylaeus signatus*.

## Notes

Conflict of interest: The authors are employed by Bayer Crop Science or BASF SE, manufacturers of crop protection products.

### Competing Interest Statement

The authors are employed by Bayer Crop Science or BASF SE, manufacturers of crop protection products.

## References

AAVM (2015). Roadmap for insect pollinator risk assessment in Australia.

Austin, A. J. and J. D. J. Gilbert (2021). “Solitary bee larvae prioritize carbohydrate over protein in parentally provided pollen.” Functional Ecology n/a(n/a).

Bosch, J. and N. Vicens (2002). “Body size as an estimator of production costs in a solitary bee.” Ecological Entomology 27(2): 129–137.

Boyle, N. K., et al. (2019). “Workshop on pesticide exposure assessment paradigm for non-Apis bees: foundation and summaries.” Environmental entomology 48(1): 4–11.

Danforth, B. N., et al. (2019). The solitary bees: biology, evolution, conservation, Princeton University Press.

Davidson, I., et al. (1986). “Biological basis for extrapolation across mammalian species.” Regulatory Toxicology and Pharmacology 6(3): 211–237.

EFSA (2009). “Risk assessment for birds and mammals.” EFSA Journal 7(12): 1438.

EFSA (2012). “Scientific Opinion on the science behind the development of a risk assessment of Plant Protection Products on bees (Apis mellifera, Bombus spp. and solitary bees).” EFSA Journal 10(5): 2668.

EFSA (2013). “EFSA Guidance Document on the risk assessment of plant protection products on bees (Apis mellifera, Bombus spp. and solitary bees).” EFSA Journal 11(7): 3295.

Erickson, H. P. (2009). “Size and shape of protein molecules at the nanometer level determined by sedimentation, gel filtration, and electron microscopy.” Biological procedures online 11(1): 32–51.

Franke, L., et al. (2021). “Results of 2-Year Ring Testing of a Semifield Study Design to Investigate Potential Impacts of Plant Protection Products on the Solitary Bees Osmia Bicornis and Osmia Cornuta and a Proposal for a Suitable Test Design.” Environmental toxicology and chemistry 40(1): 236–250.

Gallai, N., et al. (2009). “Economic valuation of the vulnerability of world agriculture confronted with pollinator decline.” Ecological economics 68(3): 810–821.

Gould, J. L. and C. G. Gould (1988). The honey bee, Scientific American Library.

Goulson, D., et al. (2015). “Bee declines driven by combined stress from parasites, pesticides, and lack of flowers.” Science 347(6229).

Grab, H., et al. (2019). “Agriculturally dominated landscapes reduce bee phylogenetic diversity and pollination services.” Science 363(6424): 282–284.

Hadley, W. (2016). ggplot2: Elegant Graphics for Data Analysis. New York, NY, Springer-Verlag

Hadley, W., et al. (2021). dplyr: A Grammar of Data Manipulation. R package version 1.0.5.

Hrassnigg, N. and K. Crailsheim (2005). “Differences in drone and worker physiology in honeybees (Apis mellifera).” Apidologie 36(2): 255–277.

Kapheim, K. M., et al. (2011). “Support for maternal manipulation of developmental nutrition in a facultatively eusocial bee, Megalopta genalis (Halictidae).” Behavioral ecology and sociobiology 65(6): 1179–1190.

Kassambara, A. (2020). Package ‘ggpubr’.

Kendall, L. K., et al. (2019). “Pollinator size and its consequences: Robust estimates of body size in pollinating insects.” Ecology and Evolution 9(4): 1702–1714.

Kyriakopoulou, K., et al. (2017). “Collection and analysis of pesticide residue data for pollen and nectar-Final Report.” EFSA Supporting Publications 14(10).

Ladurner, E., et al. (1999). “Laboratory rearing of Osmia cornuta Latreille (Hymenoptera Megachilidae) on artificial diet.” Boll. Ist. Entomol.”G. Grandi”, Univ. Bologna 53: 133–146.

Mineau, P., et al. (2001). “Reference values for comparing the acute toxicity of pesticides to birds.” Rev. Environ. Contam. Toxicol 170: 13–74.

Mineau, P., et al. (1996). “On the use of scaling factors to improve interspecies extrapolation of acute toxicity in birds.” Regulatory Toxicology and Pharmacology 24(1): 24–29.

Müller, A., et al. (2006). “Quantitative pollen requirements of solitary bees: Implications for bee conservation and the evolution of bee–flower relationships.” Biological Conservation 130(4): 604–615.

Pamminger, T. (2021). “Extrapolating acute contact bee sensitivity to insecticides based on body weight using a phylogenetically informed interspecies scaling framework.” Environmental toxicology and chemistry.

Pamminger, T., et al. (2019). “Pollen report: quantitative review of pollen crude protein concentrations offered by bee pollinated flowers in agricultural and non-agricultural landscapes.” PeerJ 7: e7394.

R (2013). “R: A language and environment for statistical computing.” RStudio (2020). RStudio: Integrated Development for R. Boston, MA, PBC.

Slowikowski, K., et al. (2016). “Package ‘ggrepel’.”

Thompson, H. M. and T. Pamminger (2019). “Are honeybees suitable surrogates for use in pesticide risk assessment for non-Apis bees?” Pest Management Science 75(10): 2549–2557.

Urban, D. H. and D. J. Urban (1986). Hazard evaluation division standard evaluation procedure: Ecological risk assessment, US Environmental Protection Agency, Office of Pesticide Programs.

USEPA (2014). “Guidance for assessing pesticide risks to bees.” Office of Chemical Safety and Pollution Prevention Office of Pesticide Programs Environmental Fate and Effects Division, Environmental Protection Agency, Washington DC; Environmental Assessment Directorate, Pest Management Regulatory Agency, Health Canada, Ottawa, CN; California Department of Pesticide Regulation.

Westrich, P. (2018). Die Wildbienen Deutschlands, Stuttgart.

Wickham, H., et al. (2019). “Welcome to the Tidyverse.” Journal of Open Source Software 4(43): 1686.

